# Structural and Biochemical Studies of Dihydrofolate Reductase from *Streptococcus pyogenes* as a Target for Antifolate Antibiotics

**DOI:** 10.1101/504357

**Authors:** Behnoush Hajian, Jolanta Krucinska, Michael Martins, Narendran G-Dayanan, Kishore Viswanathan, Sara Tavakoli, Dennis Wright

**Author notes:** Corresponding author: Dennis Wright.

## Abstract

*Streptococcus pyogenes*, a *beta*-hemolytic bacterium, causes a wide spectrum of infections in human including pharyngitis, tonsillitis, scarlet fever, rheumatic fever, and necrotizing fasciitis. Streptococcal infections can also exist as co-infection with methicillin resistant *Staphylococcus aureus* (MRSA). Trimethoprim-sulfamethoxazole (TMP-SMX) combination has been used for treatment of *S. pyogenes* and MRSA co-infection. However, resistance to TMP, an inhibitor of dihydrofolate reductase enzyme (DHFR), has challenged the efficacy of TMP-SMX combination. We explored the activity of a series of novel DHFR inhibitors against *S. pyogenes*. This study identified potent inhibitors of DHFR enzyme from *S. pyogenes* with excellent inhibitory activity against the growth of the live bacteria. We determined, for the first time, the crystal structure of *S. pyogenes* DHFR which provides structural insights into design and development of antifolate agents against this global pathogen.

## Introduction

*Streptococcus pyogenes*, or Lancefield’s group A Streptococcus (GAS), is a Gram-positive bacterium that causes a diverse spectrum of human infections. Streptococcal infections are usually mild such as pharyngitis (strep throat) and impetigo. But if the infection reaches deeper tissues, it can cause invasive infections such as necrotizing fasciitis (flesh eating disease) and streptococcal toxic shock syndrome. Superficial GAS infections can be followed by abnormal immune responses which may result in post-streptococcal sequelae including acute rheumatic fever and acute poststreptococcal glomerulonephritis.^1^ The prevalence of severe GAS infections is 18.1 million cases, with 1.78 new cases and 517 000 deaths each year. The past decade has witnessed a global resurgence of streptococcal diseases such as skin and soft tissue infections and scarlet fever.^2,3^

Although, *S. pyogenes* in general remains susceptible to most classes of antibiotics, treatment of streptococcal infections is challenged by the rising tide of antimicrobial resistance (AMR).^4–7^ Over the past two decades, there has been an increasing rate of macrolide resistance among *S. pyogenes* isolates in Europe and worldwide.^8–11^ There has been also a growing rate of penicillin failure mostly due to lack of penicillin permeation into the infected tissues and co-infection of *S. pyogenes* with beta-lactamase producing bacteria such as *Staphylococcus aureus*.^4^

Trimethoprim-sulfamethoxazole combination (SXT), one of the most widely used and cheapest antibacterials in the world, is currently suggested as a valuable option for treatment of skin and soft tissue coinfections with *S. pyogenes* and methicillin-resistant *S. aureus* (MRSA), when penicillin treatment fails.^12^ However, the emergence of trimethoprim (TMP) resistance in *S. pyogenes* isolates have challenged the efficacy of SXT.^13,14^ TMP is an inhibitor of dihydrofolate reductase (DHFR), one of the key enzymes in the folate biosynthetic pathway. The folate pathway is essential in the synthesis of reduced folates, the one-carbon donors required for the production of deoxythymidine monophosphate (dTMP), purine nucleotides, methionine and histidine (**Figure 1**). DHFR catalyzes the reduction of DHF to THF using NADPH as an electron donor. Due to its pivotal role in regulating cellular levels of THF and its derivatives, DHFR has served as an attractive target for many anticancer, antibacterial, and antiprotozoal drugs.^15,16^

**Figure 1.**
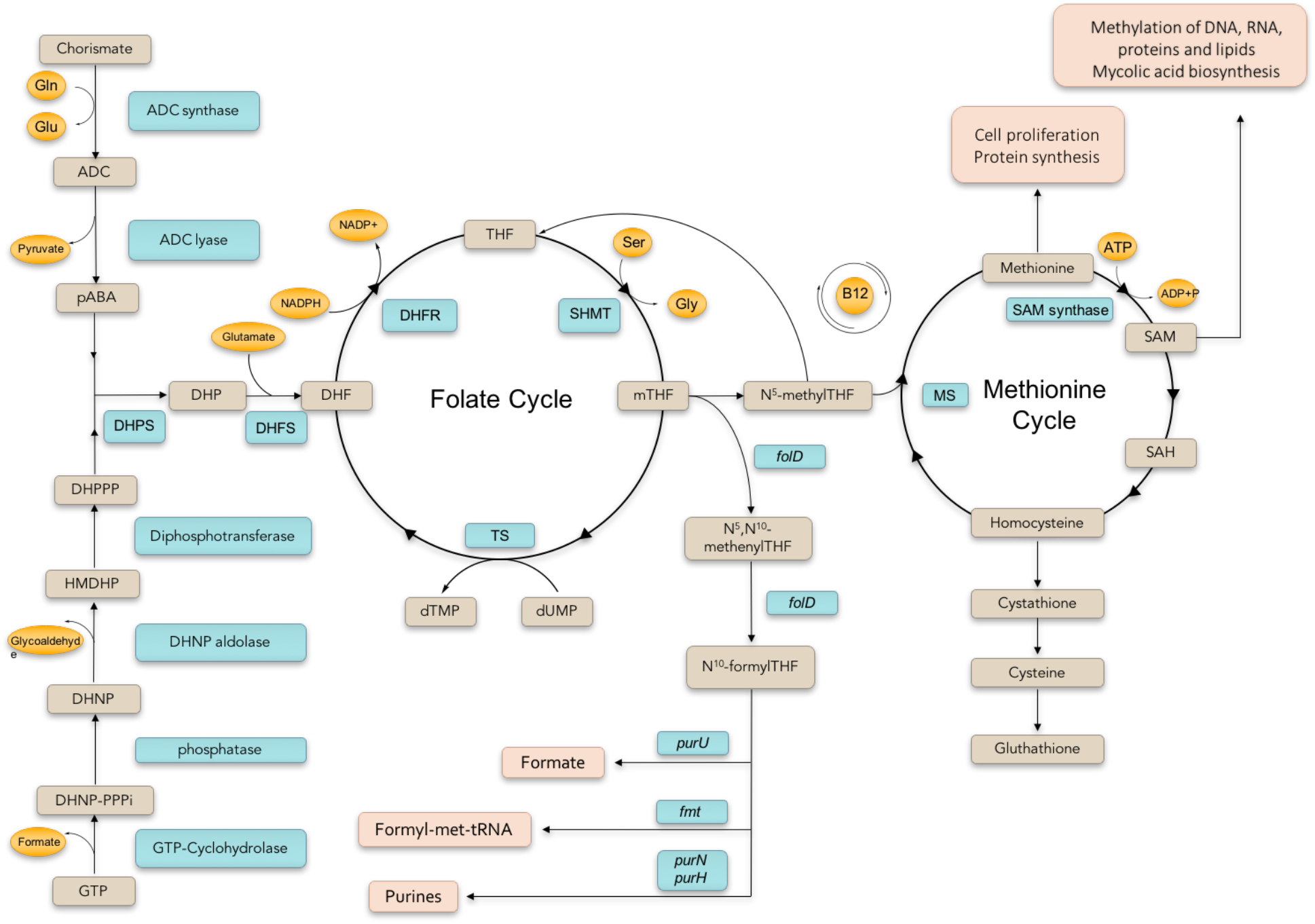
Bacterial folate cycle and related pathways. **ADC**: aminodeoxy chorismate, **PABA**: para-aminobenzoic acid, **DHPPP**: 7,8-dihydropterin pyrophosphate, **HMDHP**: 6-hydroxymethyl-7,8-dihydropterin, **DHNP**: 7,8-dihydroneopterine, **DHNP-PPPi**: 7,8-dihydroneopterine triphosphate, **DHP**: 7,8-dihydropteroate, **DHF**: 7,8-dihydrofolate, **THF**: 5,6,7,8-tetrahydrofolate, **DHFR**: dihydrofolate reductase, **SHMT**: serine hydroxymethyltransferase, **mTHF**: N^5^,N^10^-methyleneTHF, **5-mTHF**: N^5^-methylTHF, **TS**: thymidylate synthase, **MS**: methionine synthase, **SAM**: S-adenosyl methionine, **SAH**: S-adenosyl homocysteine

Herein, we report the activity of a series of propargyl-linked antifolates (PLAs) against *S. pyogenes* and DHFR enzyme from this pathogen (SpDHFR). Previously, we have shown that PLAs are inhibitors of TMP-susceptible and TMP-resistant MRSA isolates.^17–20^ Screening of PLAs against *S. pyogenes* identified a group of novel PLAs with pronounced antibacterial activity. Further exploration of structure activity relationship (SAR) of this group has led to the identification of potent inhibitors of SpDHFR enzyme.

Here we report for the first-time high resolution crystal structure of SpDHFR in a ternary complex with NADPH and one of the lead PLAs which provides structural insights into the design of potent and selective inhibitors. Our data strongly support the effort to explore PLAs as promising candidate for design of novel antifolates against TMP-resistant *S. pyogenes* and MRSA coinfections.

## Results

### Antimicrobial activity of PLAs against S. pyogenes

Using broth micro dilution method, a series of PLAs were tested for their inhibitory activity against the growth of *S. pyogenes* ATCC 19615 (Rosenbach), a quality control strain used in a variety of susceptibility assays. PLASs scaffold includes a 2,4-diaminopyrimidine ring (ring A) linked to an aryl or heteroaryl system (ring B and C) through a propargyl bridge. Based on the variations in the B- and C-rings, the tested PLAs were categorized into six general groups (**Table 1 and Table S1**). With only one exception (compound **34**), all the tested compounds exhibited very promising antimicrobial activity against *S. pyogenes* with MIC values below 1 µg/ml.

**Table 1.**
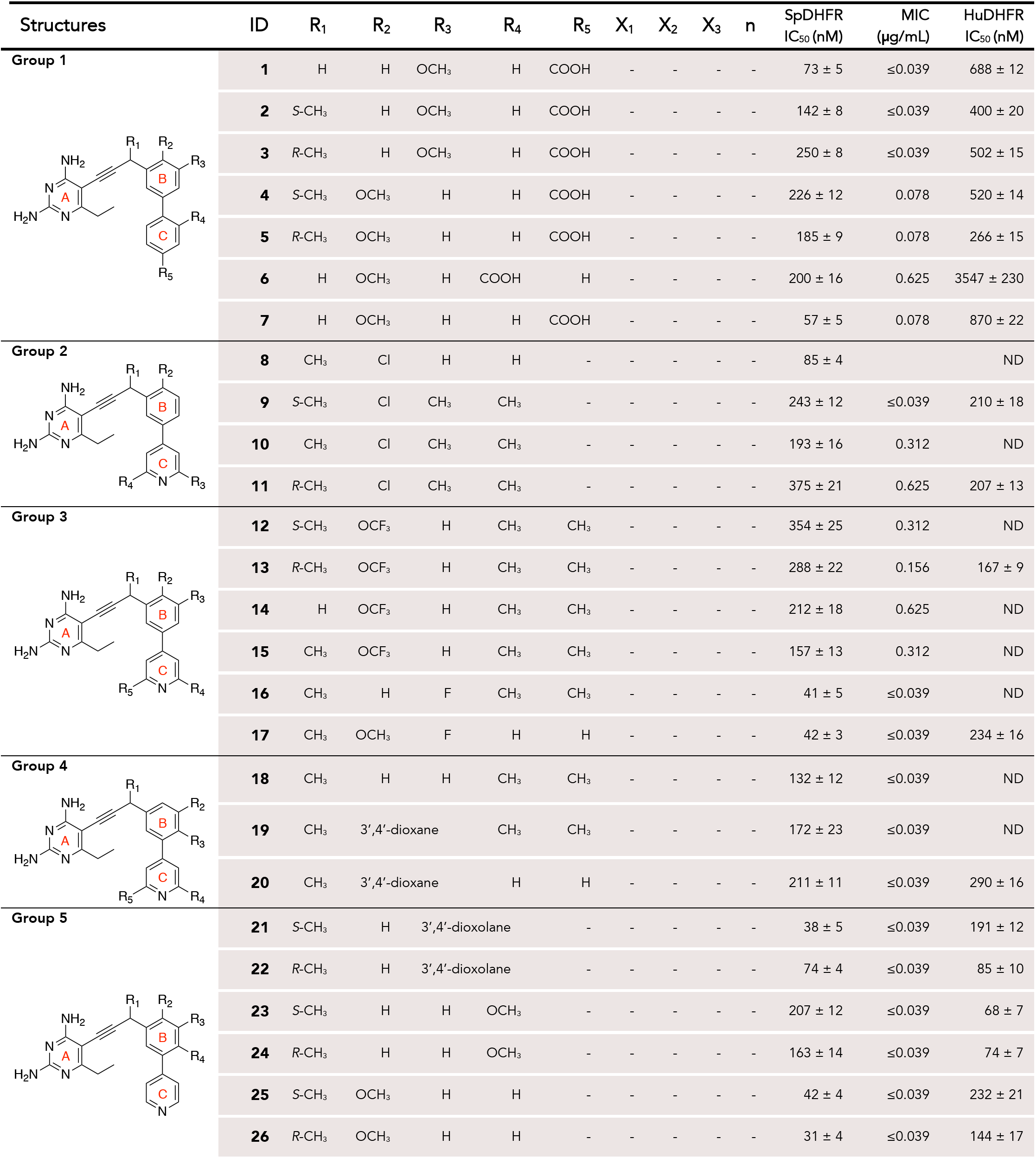
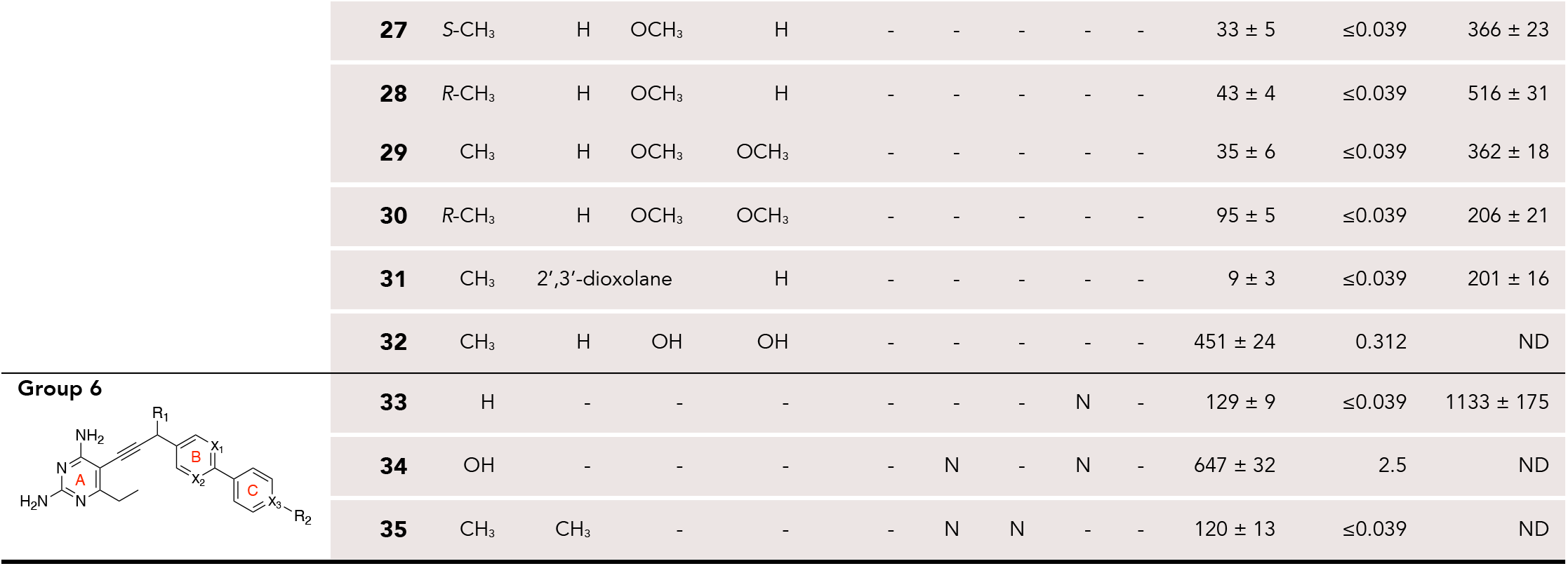
Biological activity of PLAs against *S. pyogenes* and DHFR enzymes.

| Structures | ID | R<sub>1</sub> | R<sub>2</sub> | R<sub>3</sub> | R<sub>4</sub> | R<sub>5</sub> | X<sub>1</sub> | X<sub>2</sub> | X<sub>3</sub> | n | SpDHFR<br>IC<sub>50</sub> (nM) | MIC<br>(μg/mL) | HuDHFR<br>IC<sub>50</sub> (nM) |
| --- | --- | --- | --- | --- | --- | --- | --- | --- | --- | --- | --- | --- | --- |
| **Group 1** | **1** | H | H | OCH<sub>3</sub> | H | COOH | - | - | - | - | 73 ± 5 | ≤0.039 | 688 ± 12 |
|  | **2** | S-CH<sub>3</sub> | H | OCH<sub>3</sub> | H | COOH | - | - | - | - | 142 ± 8 | ≤0.039 | 400 ± 20 |
|  | **3** | R-CH<sub>3</sub> | H | OCH<sub>3</sub> | H | COOH | - | - | - | - | 250 ± 8 | ≤0.039 | 502 ± 15 |
|  | **4** | S-CH<sub>3</sub> | OCH<sub>3</sub> | H | H | COOH | - | - | - | - | 226 ± 12 | 0.078 | 520 ± 14 |
|  | **5** | R-CH<sub>3</sub> | OCH<sub>3</sub> | H | H | COOH | - | - | - | - | 185 ± 9 | 0.078 | 266 ± 15 |
|  | **6** | H | OCH<sub>3</sub> | H | COOH | H | - | - | - | - | 200 ± 16 | 0.625 | 3547 ± 230 |
|  | **7** | H | OCH<sub>3</sub> | H | H | COOH | - | - | - | - | 57 ± 5 | 0.078 | 870 ± 22 |
| **Group 2** | **8** | CH<sub>3</sub> | Cl | H | H | - | - | - | - | - | 85 ± 4 |  | ND |
|  | **9** | S-CH<sub>3</sub> | Cl | CH<sub>3</sub> | CH<sub>3</sub> | - | - | - | - | - | 243 ± 12 | ≤0.039 | 210 ± 18 |
|  | **10** | CH<sub>3</sub> | Cl | CH<sub>3</sub> | CH<sub>3</sub> | - | - | - | - | - | 193 ± 16 | 0.312 | ND |
|  | **11** | R-CH<sub>3</sub> | Cl | CH<sub>3</sub> | CH<sub>3</sub> | - | - | - | - | - | 375 ± 21 | 0.625 | 207 ± 13 |
| **Group 3** | **12** | S-CH<sub>3</sub> | OCF<sub>3</sub> | H | CH<sub>3</sub> | CH<sub>3</sub> | - | - | - | - | 354 ± 25 | 0.312 | ND |
|  | **13** | R-CH<sub>3</sub> | OCF<sub>3</sub> | H | CH<sub>3</sub> | CH<sub>3</sub> | - | - | - | - | 288 ± 22 | 0.156 | 167 ± 9 |
|  | **14** | H | OCF<sub>3</sub> | H | CH<sub>3</sub> | CH<sub>3</sub> | - | - | - | - | 212 ± 18 | 0.625 | ND |
|  | **15** | CH<sub>3</sub> | OCF<sub>3</sub> | H | CH<sub>3</sub> | CH<sub>3</sub> | - | - | - | - | 157 ± 13 | 0.312 | ND |
|  | **16** | CH<sub>3</sub> | H | F | CH<sub>3</sub> | CH<sub>3</sub> | - | - | - | - | 41 ± 5 | ≤0.039 | ND |
|  | **17** | CH<sub>3</sub> | OCH<sub>3</sub> | F | H | H | - | - | - | - | 42 ± 3 | ≤0.039 | 234 ± 16 |
| **Group 4** | **18** | CH<sub>3</sub> | H | H | CH<sub>3</sub> | CH<sub>3</sub> | - | - | - | - | 132 ± 12 | ≤0.039 | ND |
|  | **19** | CH<sub>3</sub> | 3',4'-dioxane |  | CH<sub>3</sub> | CH<sub>3</sub> | - | - | - | - | 172 ± 23 | ≤0.039 | ND |
|  | **20** | CH<sub>3</sub> | 3',4'-dioxane |  | H | H | - | - | - | - | 211 ± 11 | ≤0.039 | 290 ± 16 |
| **Group 5** | **21** | S-CH<sub>3</sub> | H | 3',4'-dioxolane |  | - | - | - | - | - | 38 ± 5 | ≤0.039 | 191 ± 12 |
|  | **22** | R-CH<sub>3</sub> | H | 3',4'-dioxolane |  | - | - | - | - | - | 74 ± 4 | ≤0.039 | 85 ± 10 |
|  | **23** | S-CH<sub>3</sub> | H | H | OCH<sub>3</sub> | - | - | - | - | - | 207 ± 12 | ≤0.039 | 68 ± 7 |
|  | **24** | R-CH<sub>3</sub> | H | H | OCH<sub>3</sub> | - | - | - | - | - | 163 ± 14 | ≤0.039 | 74 ± 7 |
|  | **25** | S-CH<sub>3</sub> | OCH<sub>3</sub> | H | H | - | - | - | - | - | 42 ± 4 | ≤0.039 | 232 ± 21 |
|  | **26** | R-CH<sub>3</sub> | OCH<sub>3</sub> | H | H | - | - | - | - | - | 31 ± 4 | ≤0.039 | 144 ± 17 |
|  | **27** | S-CH<sub>3</sub> | H | OCH<sub>3</sub> | H | - | - | - | - | - | 33 ± 5 | ≤0.039 | 366 ± 23 |
|  | **28** | R-CH<sub>3</sub> | H | OCH<sub>3</sub> | H | - | - | - | - | - | 43 ± 4 | ≤0.039 | 516 ± 31 |
|  | **29** | CH<sub>3</sub> | H | OCH<sub>3</sub> | OCH<sub>3</sub> | - | - | - | - | - | 35 ± 6 | ≤0.039 | 362 ± 18 |
|  | **30** | R-CH<sub>3</sub> | H | OCH<sub>3</sub> | OCH<sub>3</sub> | - | - | - | - | - | 95 ± 5 | ≤0.039 | 206 ± 21 |
|  | **31** | CH<sub>3</sub> | 2',3'-dioxolane |  | H | - | - | - | - | - | 9 ± 3 | ≤0.039 | 201 ± 16 |
|  | **32** | CH<sub>3</sub> | H | OH | OH | - | - | - | - | - | 451 ± 24 | 0.312 | ND |
| **Group 6** | **33** | H | - | - | - | - | - | - | N | - | 129 ± 9 | ≤0.039 | 1133 ± 175 |
|  | **34** | OH | - | - | - | - | N | - | N | - | 647 ± 32 | 2.5 | ND |
|  | **35** | CH<sub>3</sub> | CH<sub>3</sub> | - | - | - | N | N | - | - | 120 ± 13 | ≤0.039 | ND |

### Inhibition of S. pyogenes DHFR enzyme by PLAs

To test whether the inhibitory activity of PLAs against *S. pyogenes* cells is mediated through the inhibition of DHFR enzyme, we evaluated the inhibitory effect of the compounds against the purified enzyme. The half maximal inhibitory concentrations (IC_50_ values) are shown in **Table 1**. Although the potency of the compounds varies, in general there is a correlation between the enzyme and cell growth inhibition. SAR analysis of each group of the tested compounds elucidated key structural features that affect the potency of PLAs against SpDHFR.

Group one is characterized by PLACOOH compounds which feature a carboxylic acid substitution on the C-ring. The IC_50_ values observed in this category varies between 50 to 250 nM. Within this group, the presence of propargylic methyl correlates with a decrease in the potency of the compounds. As seen with compounds **1** and **7**, the absence of propargylic methyl leads to improved inhibitory activity against SpDHFR (IC_50_ values of 73 and 57 nM, respectively). In addition, moving the carboxylate from the *para* (in compound **7**) to the *ortho* position (in compound **6**) compromised the inhibitory effect by almost four-folds (IC_50_ values of 57 and 200 nM, respectively). Furthermore, the stereoisomers of *S* configuration (compounds **2** and **4**) exhibit similar activity with *R* isomers (compounds **3** and **5**). The lack of apparent stereospecificity is continuously evident across all the other tested PLAs.

Compounds in group two feature a pyrimidine C-ring and a chlorine substitution at 2’ position on the B-ring. Here, compound **8** is the most potent congener with IC_50_ value of 85 nM. Introduction of two methyl groups on the C-ring (compound **10**) leads to a modest decrease in inhibitory affinity (IC_50_ value of 193 nM).

Based on the same concept, group three represents fluorine atoms substitution on the B-ring. IC_50_ values for this group range from 40 to 350 nM. Not surprisingly, the presence of propargylic methyl and/or methyl substitutions on the C-ring reflects the same trend as seen with chlorine atom derivatives. Next we evaluated the effect of heterocyclic substitution on the B-ring by introducing a dioxane and dioxalane moieties on the B-ring (Group four and five).

Interestingly, the presence of 3’,4’-dioxane (compounds **19** and **20**) has minimal effect on the potency of PLAs (IC_50_ values of 132 and 172 nM). By contrast, 3’-4’-dioxalane substitution (compound **21** and **22**) resulted in enhance inhibitory activity by as much as five-fold.

These observations prompted us to investigate different B-ring modifications in the contest of substituted and unsubstituted B-ring (group five). Moving the dioxolane substitution from 3′,4′ to 2′,3′ position yielded compound **31**, superior in both potency and selectivity, with IC_50_ value of 9 nM. These structural analysis highlights the importance of a simple change around the B-ring at 2′, 3′, and 4′ position on the potency and affinity of these compounds against SpDHFR.

Group six represent three compounds with *para* C-ring and various nitrogen substitutions on B- and C-rings. These compounds have moderate inhibitory activity with IC_50_ values above 100 nM. It is notable that replacement of the propargylic methyl with a hydroxyl moiety (compound **34**) is not tolerated (IC_50_ value of ∼650 nM). Selectivity over the human form of DHFR, to ensure low toxicity, is an important parameter to consider. We have measured the inhibitory effect of select compounds against the human DHFR enzyme (HuDHFR) and reported it as IC_50_ values (**Table 1**).

In summary, compounds lacking propargylic substitution (compounds **1** and **7**) and/or those without bulky substitutions on the C-ring yielded more selective compounds. The main structural variations that drive the potency and selectivity of PLAs against SpDHFR appear to be the simplified propargyl, 2′, and 3′ substitution on the B-ring as well as meta-substitution on the C-ring.

### Enzyme Structural Analysis

To investigate drug-target interactions, we have successfully crystalized SpDHFR in complex with compound **3**. The X-ray crystal structure of SpDHFR bound to cofactor NADPH and the ligand was determined at 2.2 Å with the final R-factor of 0.19 and R_free_ of 0.25 (**Table S1**). The crystal belongs to the orthorhombic space group P2_1_2_1_2_1_ with four molecules in the asymmetric unit. Despite low sequence homology with DHFR enzymes (**Figure 2**), SpDHFR structure exhibit the same general fold composed of a central b-sheet and four flanking α-helices (**Figure 3 A and B**). SpDHFR is composed of eight parallel β-strands (β1, β2, β3, β4, β5, β6, β7, and β9), two anti-parallel β-strands (β8 and β10), and four α-helices (α1, α2, α3, and α4). Full density for both NADPH and compound **3** was also observed (**Figure 4A**).

**Figure 2.**
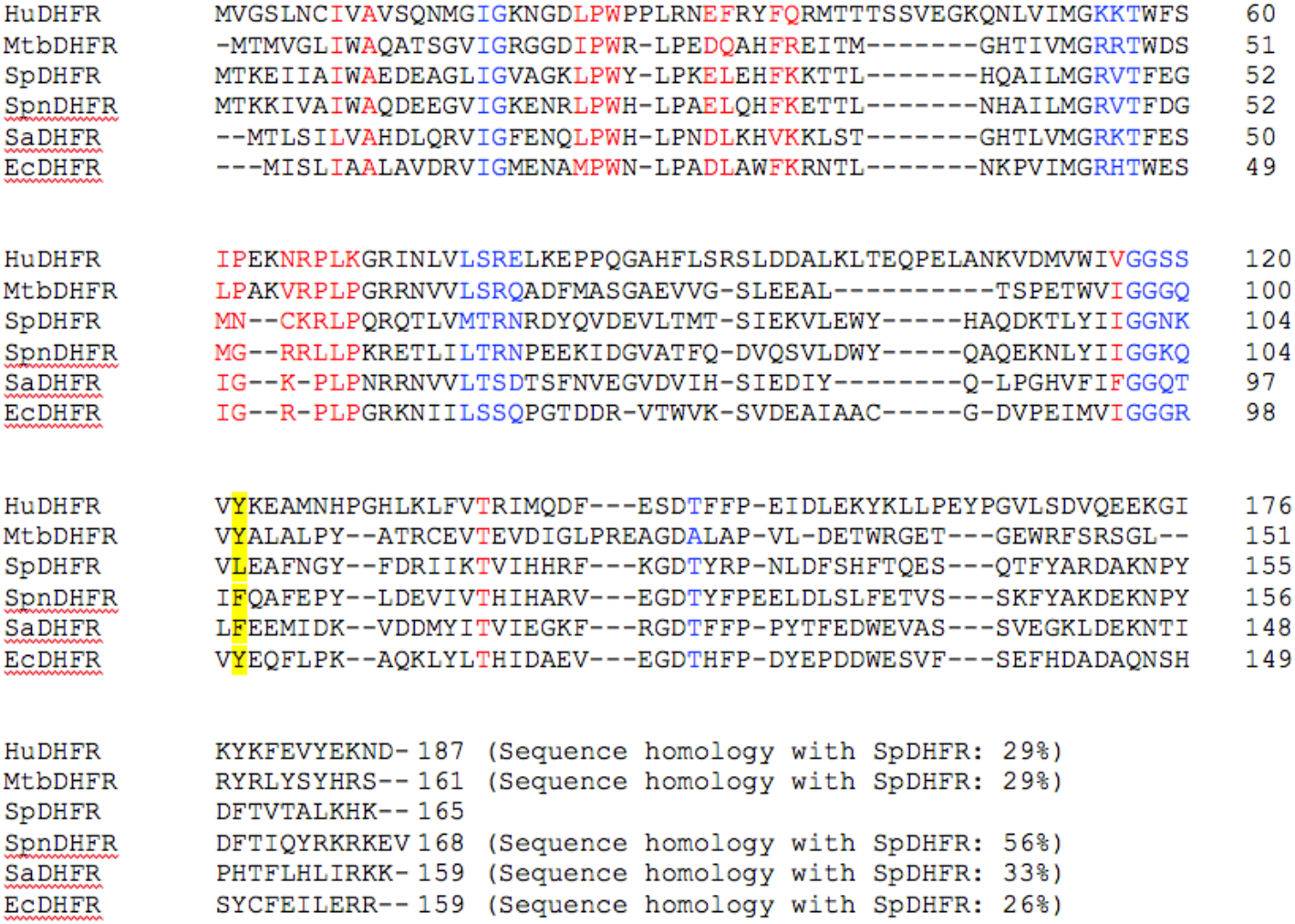
Sequence alignment of SpDHFR with other known DHFR enzymes. Folate binding site residues are shown in red. Cofactor binding site residues are shown in blue. **Hu**: *Homo sapiens*, **Mtb**: *Mycobacterium tuberculosis*, **Spn**: *Streptococcus pneumoniae*, **Sa**: *Staphylococcus aureus*, and Ec: *Escherichia coli*

**Figure 3.**
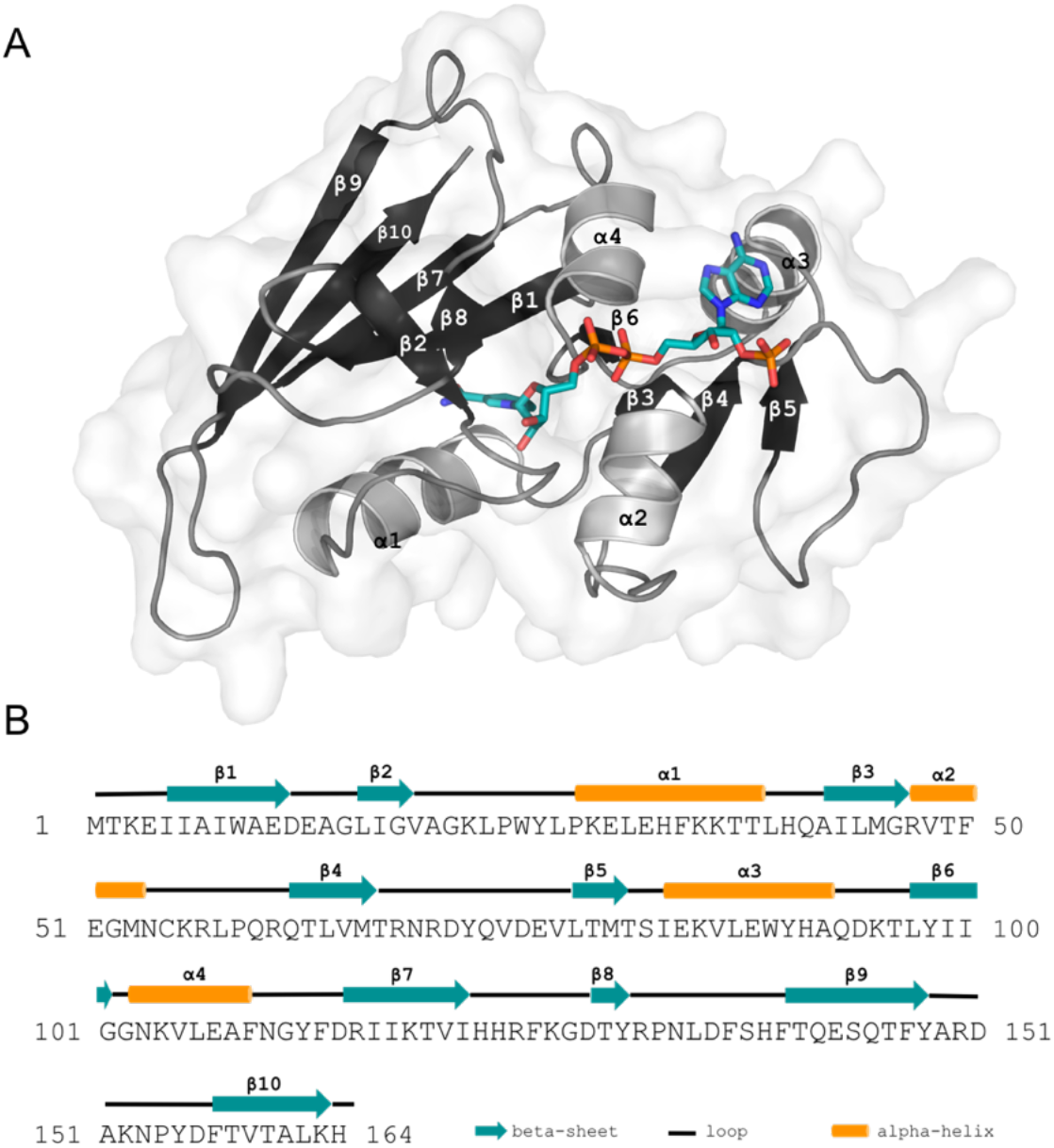
Structure of SpDHFR and assignment of secondary structures. A) Schematic ribbon diagram of the overall fold of SpDHFR. NADPH is shown using a stick model. B) Assignment of secondary structures for SpDHFR sequence.

**Figure 4.**
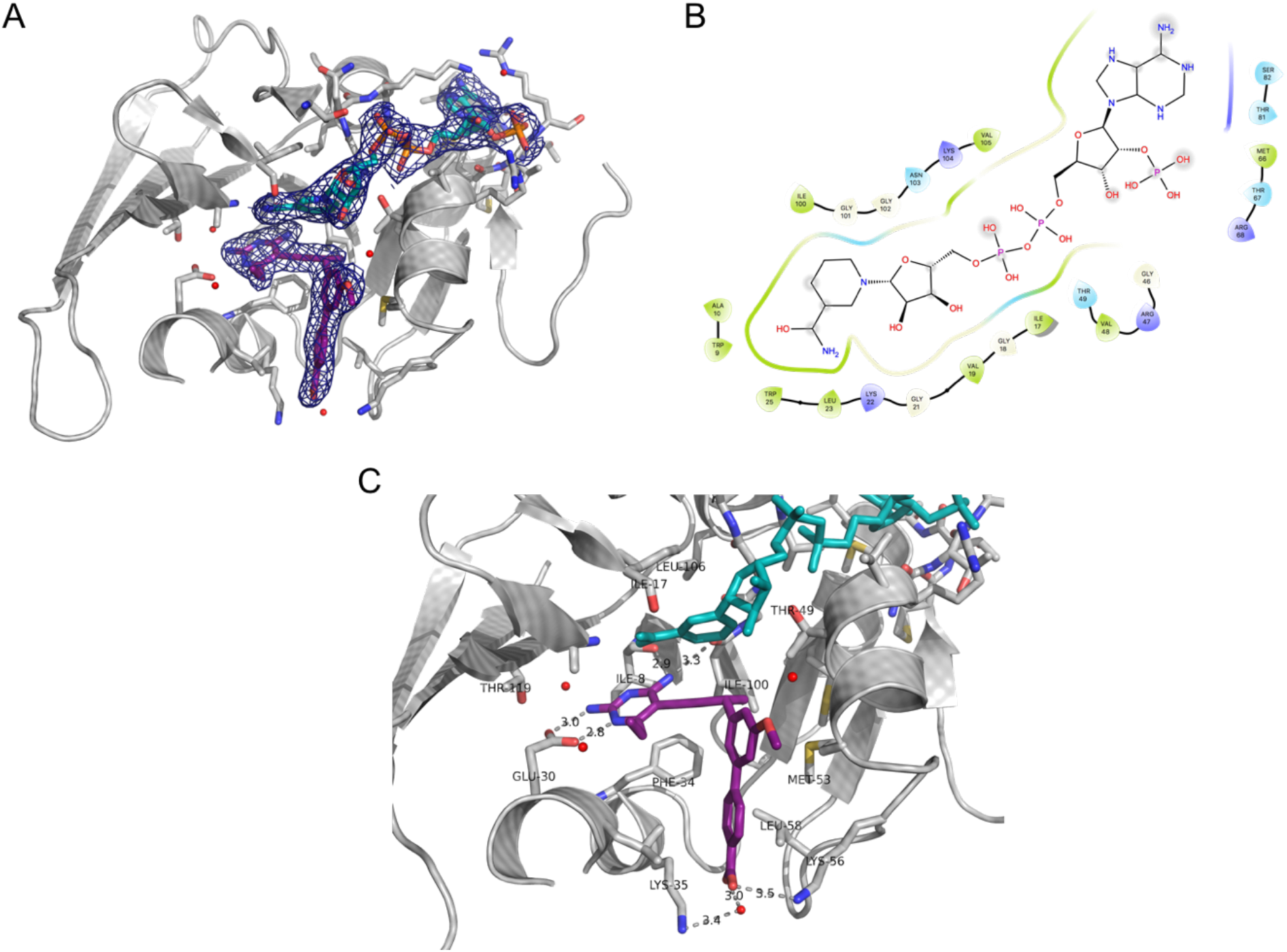
A) Structure of SpDHFR in complex with NADPH and compound 3, B) 2D diagram of NADPH interactions with the binding site residues, C) A detailed view of SpDHFR interactions with compound 3

### NADPH binding site

The NADPH molecule is bound to SpDHFR in an extended conformation with the nicotinamide ring inserted into a cleft formed by β1, β2, and β8-strands (**Figures 3A and 4A**). NADPH is anchored into the cofactor binding pocket through extensive interactions with the active site residues (**Figure 4B**). The amide group of the nicotinamide ring forms three hydrogen bonds with the backbone of Ala10 and Ile17. The nicotinamide ribose contacts Val19, Gly21, and Lys22. The pyrophosphate moiety interacts extensively with the residues from α2 and α4 helices (Val48, Thr49, Asn103, and Lys104). The O2′-phosphate of adenosyl ribose forms four hydrogen bonds with Arg47, Thr67, and Arg68. The adenine group contacts the protein through interactions with Thr81 and Ser82 and stacking against Met66.

### Folate binding site

The 2,4-diaminopyrimidine ring of compound **3** is deeply positioned into the hydrophobic binding pocket and forms two hydrogen bonds with Glu30 and another hydrogen bond with the backbone carboxylate of Ile8 (**Figures 4A and 4C**). Glu30 is highly conserved and critical for the catalytic activity of DHFR. In some species, this residue is replaced by Asp which provides similar interactions (**Figure 2**). There are also water-mediated interactions between the 2,4-diaminopyrimidine ring of compound **3** and the residues in the binding pocket and p-p stacking with Phe34. This arrangement around the diaminopyrimidine ring is highly conserved among the known DHFR structures from various species. The biaryl moiety of compound **3** makes hydrophobic interactions with Il100, Thr49, Phe50, Met53, and Leu58. The carboxylate of compound **3** has a weak electrostatic interaction with Lys 56 (3.5 Å) and a water-mediated interaction with Lys35.

### Comparison of DHFR enzymes from S. pyogenes and S. aureus

Despite low sequence homology (33%), the general fold of the two protein is very similar. A direct comparison of the active site residues in SpDHFR structure with the previously reported structure of DHFR enzyme from *S. aureus* (PDB ID: 4Q67)^21^ revealed highly conserved binding pocket (**Figure 5A**) which should allow for the design of dual inhibitors of both enzymes.

**Figure 5.**
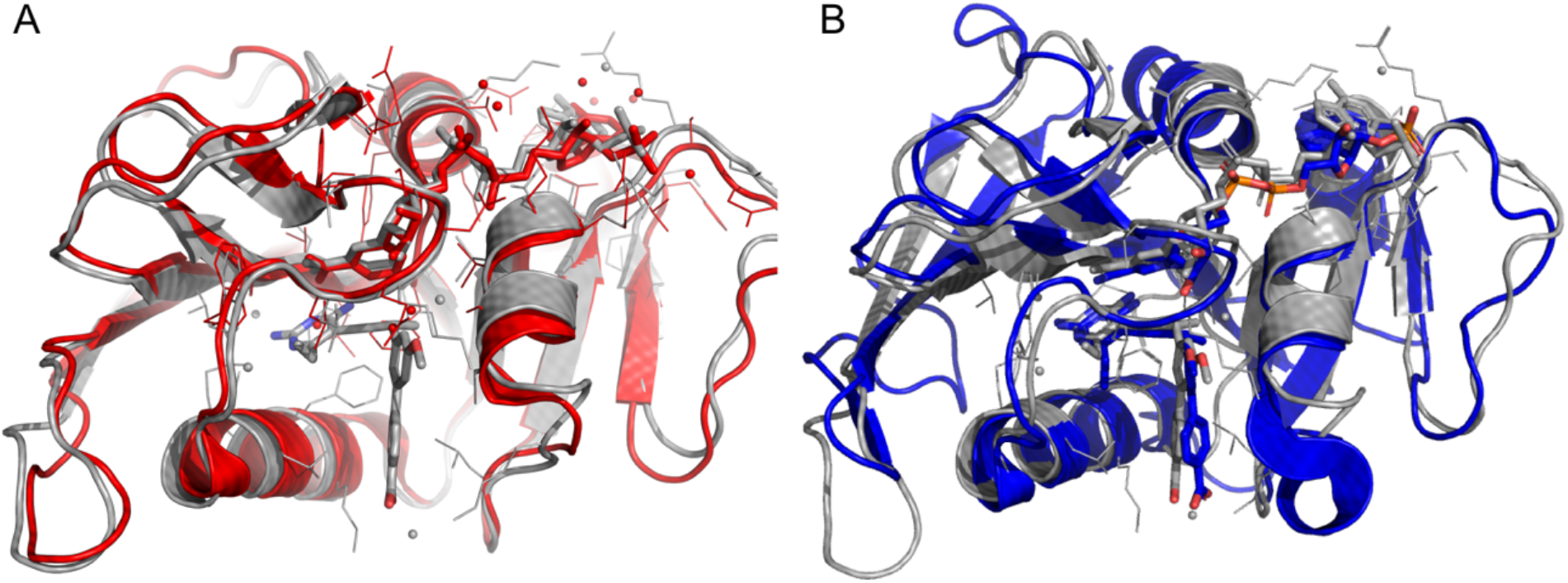
A) Superposition of SpDHFR (gray) and SaDHFR (red), B) Superposition of SpDHFR (grey) and HuDHFR (blue)

### Selectivity and Comparison with HuDHFR

SpDHFR contains 165 amino acids compared with 187 residues in the HuDHFR. Despite the larger size of the human protein and low sequence homology (29%), the overall folding of the two proteins is very similar. By superimposing SpDHFR structure with human DHFR structure bound to the same ligand^22^ (**Figure 5B**), it becomes evident that the interactions of the 2,4-diaminopyrimidine ring that anchors compound **3** in the active site are almost identical.

Despite the similarities, there are several structural variations that can be exploited for design of selective inhibitors. For example, replacement of Phe31 in HuDHFR by corresponding residue Leu31 in SpDHFR provides greater van der Waals interactions in SpDHFR while the larger group likely contribute to destabilizing interactions in HuDHFR. Another notable difference is Pro61 in HuDHFR which corresponds to Asn54 in SpDHFR. Inhibitors with polar substitutions that can advantage from contacts with Asn54 may bind selectively to SpDHFR. An analog of compound **3** with a hydrogen bond donor on the C-ring or a hydrophilic group added to the B-ring may provide an excellent chemical space to explore. Still, a consideration should be given to the size, position, and character of these additional moieties as seen from our SAR analysis.

## Conclusion

The results presented in this work, provide insights into the structure-based design of inhibitors of DHFR in *S. pyogenes*. DHFR inhibitors have been widely used as effective anticancer and antibacterial agents.^15,23^ However, resistance to TMP, one of the most effective DHFR inhibitors, has challenged the efficacy of this drug as an antibacterial agent. TMP, often in combination with SMX, is used for treatment of skin, urinary tract, and enteric infections caused by Gram-positive and Gram-negative bacteria.^24–26^ *S. pyogenes* causes million cases of human infections every year and its capacity to develop invasive infections emphasizes the need for global control of streptococcal infections. Common co-infection of *S. pyogenes* and MRSA and difficulties in distinguishing between the two, highlights the importance of developing therapeutics with activity against both pathogen and ideally TMP-resistant isolates. Previously, we have reported a series of DHFR inhibitors with activity against TMP-sensitive and TMP-resistant MRSA isolates.^17–20^ Here, we have shown that these compounds maintain their inhibitory activity against DHFR enzyme from *S. pyogenes* and the growth of live bacteria. SAR analysis of the tested compounds along with determining the structure of SpDHFR in complex with one of the lead compounds provide key structural information required for optimization of dual inhibitors of both pathogen.

## Methods

### Antimicrobial agents

The synthesis and characterization of PLAs have been described in several publications. Trimethoprim (TMP) was purchased from Sigma Chemical Co., St. All the compounds were dissolved in 100% dimethyl sulfoxide (DMSO) prior to use.

### Bacterial isolate

S. pyogenes ATCC 19615 was purchased from the American Type Culture Collection (Manassas, VA). The organism was grown in Todd Hewitt broth.

### In vitro susceptibility testing

The assay was performed in Isosensitest broth (Oxoid) supplemented with 5% defibrinated sheep blood (ThermoFisher). Minimum inhibitory concentrations (MIC) were determined by broth microdilution method based on CLSI guideline using a final inoculum of 5 × 10^5^ CFU/ml. The antimicrobial agents were prepared at 40 µg/ml and were dispensed using serial two-fold dilution. The MIC was defined as the lowest concentration of antimicrobial agent yielding no visible growth after monitoring cell turbidity following an incubation period of 24–36 hours at 37°C.

### Transformation, expression, and purification of SpDHFR

Recombinant pET-24a(+) plasmid harboring the *folA* gene encoding SpDHFR was constructed by GenScript. BL21(DE3) competent *E. Coli* cells (New England BioLabs) were transformed with the recombinant plasmids. Transformed cells were grown in LB medium supplemented with 30 µg/mL kanamycin at 37°C until OD_600_ reached 0.6–0.7. The cells were induced with 1 mM IPTG for 20 hours at 20°C and spun down at 8000 rpm for 15 minutes. Each gram of wet cell pellet was resuspended in 5 ml of lysis buffer (25 mM Tris pH 8.0, 0.4 M KCl, 5 mM imidazole, 5 mM BME, 5% glycerol, 200 µg/ml lysozyme, 1 mM DNase I). The cell suspension was incubated for 30–60 minutes at 4°C with gentle rotation followed by sonication until a homogenous lysate was obtained. The lysate was centrifuged at 18,000 rpm for 30 minutes and supernatant was collected and filtered through 0.22 µm filter. The SpDHFR construct did not contain histidine tag and were purified over methotrexate-agarose column pre-equilibrated with 4 CV of equilibration buffer (20mM Tris-HCl pH 7.5, 50 mM KCl, 2 mM DTT, 0.1 mM EDTA and 15% glycerol). The column was washed with 3 CV of wash buffer (20 mM Tris-HCl pH 7.5, 500 mM KCl, 2 mM DTT, 0.1 mM EDTA and 15% glycerol). The protein was eluted with 3 CV of elution buffer (equilibration buffer pH 8.5 + 2mM DHF). Fractions containing SpDHFR protein were collected, concentrated and loaded onto a Hi-Prep 26/60 Sephacryl s-200 HR prepacked gel filtration/size exclusion column pre-equilibrated with 1 CV of final buffer (25 mM Tris pH 8.0, 50 mM KCl, 0.1 mM EDTA, 2 mM DTT and 15% glycerol). The column was washed with another 1 CV of final buffer and protein elution was monitored with AKTA UV/vis diode array spectrophotometer at 280 nm. Fractions containing pure enzyme were pooled, concentrated at 10 mg/ml and flash frozen in liquid nitrogen and stored at −80°C.

### Enzyme inhibition assay

The DHFR activity of SpDHFR was measured in 500 µl of assay buffer containing 20 mM TES, 50 mM KCl, 0.5 mM EDTA, 10 mM 2-mercaptoethanol (BME) and 1 mg/ml bovine serum albumin (BSA) with various concentrations of NADPH and DHF ranging from 0 to 100 µM. Assay was started by adding DHF and monitoring NADPH oxidation at 340 nm. All measurements were performed at room temperature and in triplicates. Initial velocity data were fitted with the Michaelis-Menten equation using Graphpad Prism 7.0 software. The DHFR activity inhibition assays and IC_50_ determination were performed in the same assay buffer with 100 µM NADPH and 100 µM DHF. Inhibitors, dissolved in 100% DMSO, were added to the mixture and incubated for 5 minutes before the addition of DHF. Average IC_50_ values and standard deviations were measured in triplicate.

### Crystallization of SpDHFR

All the crystallization trials were performed by hanging drop vapor diffusion method and using EasyXtal 15-well plates (Qiagen). SpDHFR was mixed with 10 mM NADPH and 2mM ligand, incubated on ice for two hours and concentrated to 14 mg/mL. 2 µl of this solution was mixed with 2 µl of crystallization solution containing 50–150 mM sodium cacodylate pH 6–7, 200 mM magnesium acetate, and 20–30% of polyethylene glycol 3350. Small polygonal crystals grew within 2–3 weeks at 4°C. Crystals were flash frozen in the mother liquor supplemented with 20% glycerol.

### Data collection and structure determination

X-ray data were collected at National Synchrotron Light Source II (NSLS II) at Brookhaven National Laboratory. Data were integrated using iMOSFLM and scaled and merged using AIMLESS from CCP4i2 suite. Molecular replacement was performed using Phaser and previously reported structure of DHFR from S. pneumoniae sharing ∼50% sequence identity with SpDHFR. The structure was refined using Coot and Phenix softwares.

