## Supplementary material for "Structural and Biochemical Studies of Dihydrofolate Reductase from *Streptococcus pyogenes* as a Target for Antifolate Antibiotics": Figure S1 and Table S1

**Figure S1. Chemical Structure of the compounds tested against *S. pyogenes* and SpDHFR**

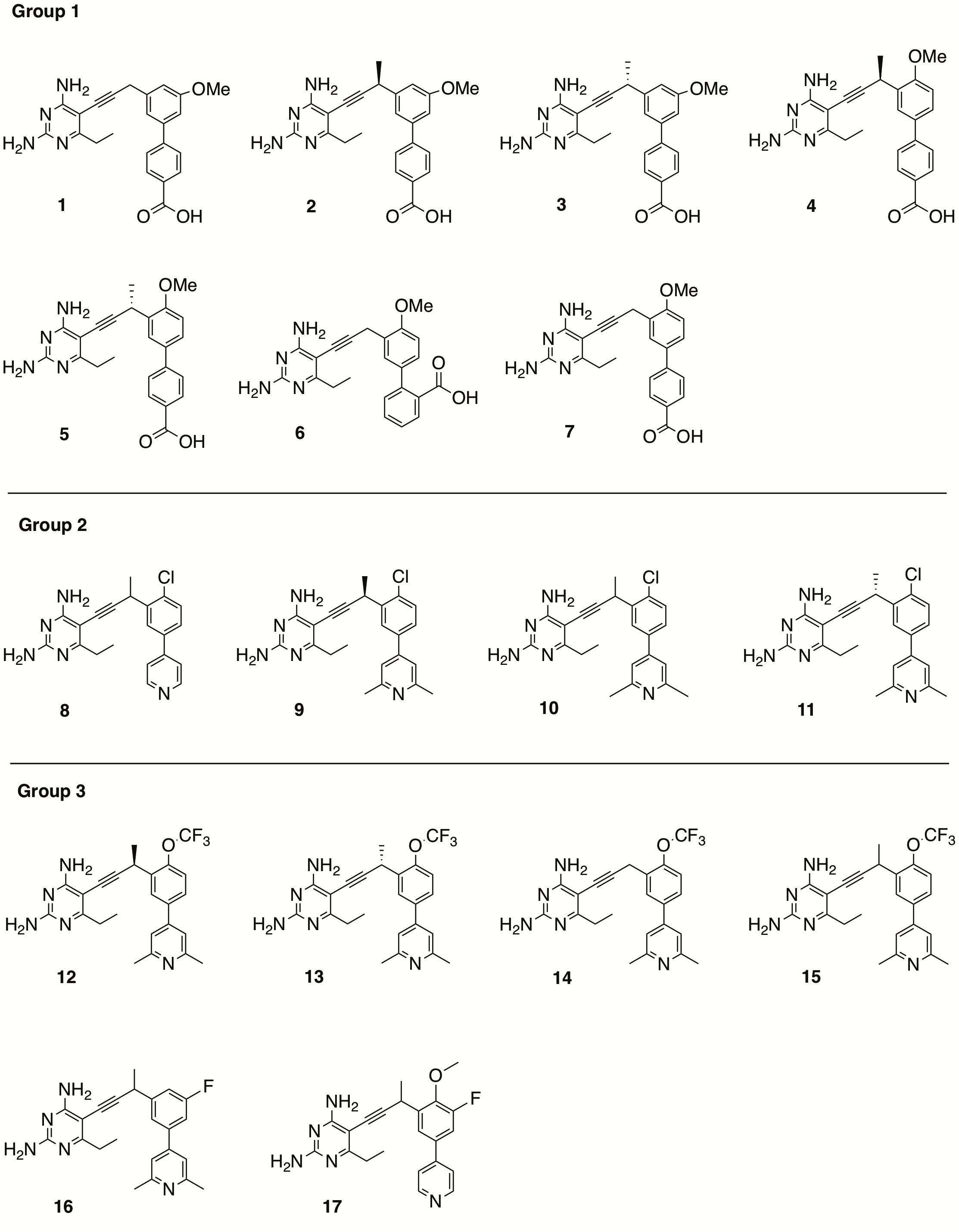

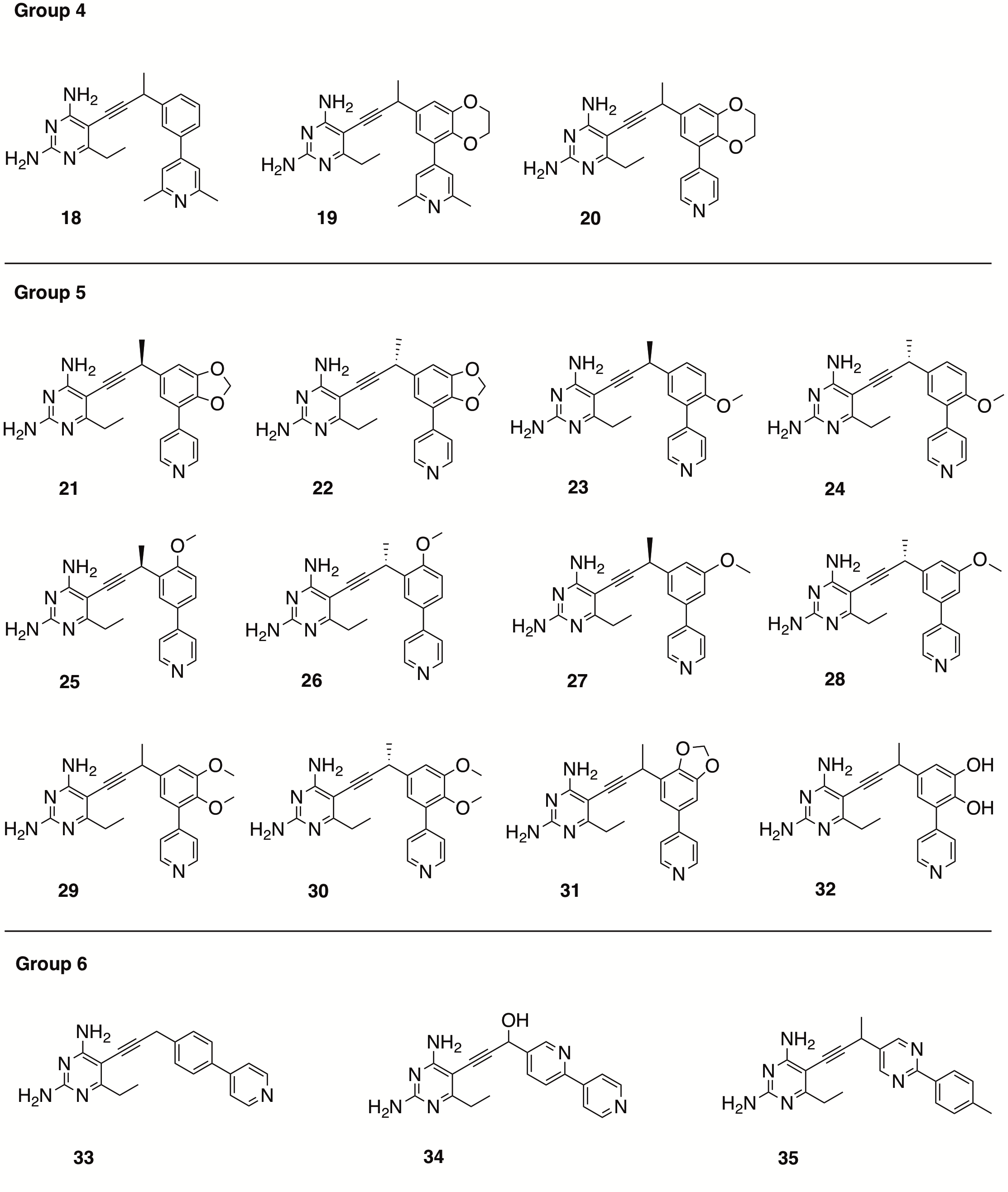

**Table S1. Crystallography data collection and refinement statistics**

| Complex | MtbDHFR:NADPH:1172 |
| --- | --- |
| PDB ID | XXXX |
| Data collection statistics | |
| Beamline | NSLS-II (FMX) |
| Space group | P2_1_2_1_2_1_ |
| Cell dimensions | |
| a, b, c (Å) | 77.69, 104.13, 119.32 |
| α, β, γ (°) | 90, 90, 90 |
| Resolution (Å) | 78.5-2.2 |
| Highest resolution shell (Å) | 2.24-2.2 |
| Completeness% | 100 (100) |
| R_sym_ |  |
| R_pim_ | 0.05 (0.27) |
| R_meas_ | 0.04 (0.28) |
| R_merge_ | 0.16 (0.89) |
| Redundancy | 2 |
| Mean I/σ(I) | 9 (3.5) |
| Refinement statistics | |
| Resolution (Å) | 78.5-2.2 |
| No. of reflections | 49689 |
| R_factor_/R_free_ | 0.19/0.25 |
| No. of atoms (protein, ligand, solvent) | 6365 |
| rmsd bond length (Å) | 0.008 |
| rmsd bond angle (°) | 1.2 |
| Ramachandran plot analysis (%) | |
| Most favored | 94.7 |
| Outliers | 0.3 |
